# Functional regeneration of the Rdl-A296S mutation in *Anopheles funestus* decouples insecticide resistance from genetic background and isolates its life-history costs

**DOI:** 10.64898/2026.11.19.767793

**Authors:** Charlotte M. Quinn, Holly Humphreys, Odette N. Zongo, Charles S. Wondji, Tony Nolan

## Abstract

Resolving the genetic basis of insecticide resistance in *Anopheles funestus* has been limited both by the historic difficulty of germline transformation in this species and by the confounding genetic and metabolic background present in field populations. The A296S “resistance-to-dieldrin” (*Rdl*) mutation provides an informative proof-of-principle case because it has persisted at high frequency in African vector populations long after the withdrawal of dieldrin. Here, we used a marker-less CRISPR/Cas9 homology-directed repair strategy to introduce the A296S mutation in isolation into the insecticide-susceptible FANG strain of *An. funestus*. WHO tube bioassays were used to quantify resistance across different dieldrin exposure regimes and to assess cross-resistance to other insecticides, while larval development and adult longevity assays were used to evaluate life-history effects. A296S alone was sufficient to confer strong dieldrin resistance. Resistance was context-dependent: homozygous resistance remained robust at higher exposures, whereas heterozygote protection declined with increasing toxic load. No operationally relevant cross-resistance was detected to DDT or pyrethroids. The mutation imposed a modest, recessive delay in larval development, with no detectable reduction in adult longevity and no clear net disadvantage across the life-history traits measured. This study provides the first in vivo functional validation of an insecticide resistance mutation in *An. funestus*. By isolating A296S on a defined susceptible background, it provides a quantitative baseline for the effect of a single target-site mutation and offers new insight into how stage-specific fitness effects may contribute to the long-term persistence of resistance alleles in natural populations.

## INTRODUCTION

Despite being preventable and curable, malaria persists as a major global health burden, with 2024 estimates placing the toll at 282 million cases and 610,000 deaths (World Health Organization 2025). Insecticide-based vector control, including long-lasting insecticidal nets (LLINs) and indoor residual spraying (IRS), has historically driven substantial reductions in transmission. However, these gains have recently stagnated, largely due to the emergence and spread of insecticide resistance (IR) in major vector species (Bhatt et al. 2015; Hemingway et al. 2016).

In sub-Saharan Africa, the transmission landscape is undergoing a significant shift: while *Anopheles gambiae* has declined in eastern and southern regions, *Anopheles funestus* has emerged as an increasingly dominant vector (Msugupakulya et al. 2023). Its highly anthropophilic and endophilic behaviour allows it to sustain intense transmission even in settings with high intervention coverage (Matowo et al. 2023; Boddé et al. 2025). Efforts to respond to this growing threat are constrained by two key limitations. First, insecticide resistance data for *An. funestus* remain sparse relative to *An. gambiae* (IR Mapper 2024). Second, *An. funestus* has historically been difficult to colonise and genetically manipulate (Ngowo et al. 2021; Odero et al. 2023).

Traditionally, identifying the genetic basis of insecticide resistance has relied on colony-based approaches, where resistance can be tracked as a Mendelian trait. This is rarely feasible for *An. funestus*, given the limited number of stable laboratory colonies (Ngowo et al. 2021; Odero et al. 2023). Field-based approaches can provide partial insight, but genotype–phenotype associations are often confounded by genetic background and co-occurring resistance mechanisms (Wondji et al. 2011; Menze et al. 2016; Weedall et al. 2020; Hardstone and Scott 2010). Similarly, introgression of candidate resistance alleles into susceptible strains is often incomplete and may fail to capture the full haplotypic context in which resistance evolved (Grigoraki et al. 2021; ffrench-Constant and Bass 2017; Clarkson et al. 2021). More recently, population genomics approaches – including genome-wide association studies (GWAS) and scans for signatures of selection – have provided powerful tools for identifying candidate resistance loci in natural populations (Gadji et al. 2024; Clarkson, Temple, and Miles 2018). These methods have greatly expanded our ability to detect genomic regions under selection, even in species such as *An. funestus* where laboratory resources are limited (Weedall et al. 2020). Despite these advances, such approaches remain correlative and often capture signals arising from linked variation or complex haplotypes, making it difficult to resolve the contribution of individual mutations (Clarkson et al. 2021; Grau-Bové et al. 2020).

Molecular surveillance programmes hold considerable promise for tracking the emergence and spread of insecticide resistance, enabling the early detection of resistance-associated variants at scale in natural populations. For these approaches to be effective, a clear understanding of the relative importance of different circulating mutations is essential. Without quantitative estimates of effect size, it remains difficult to distinguish between mutations that are causative, compensatory, or operationally relevant, limiting their usefulness for early warning and resistance management (Donnelly, Isaacs, and Weetman 2016; Clarkson, Temple, and Miles 2018; Clarkson et al. 2021).

CRISPR/Cas9 genome editing provides a route to overcome these limitations (Grigoraki et al. 2021; Williams et al. 2022). Our group has previously demonstrated efficient knock-in of transgenes in *An. funestus*, consistent with evidence that CRISPR-induced breaks in *Anopheles* can be efficiently repaired via homology-directed repair (HDR) (Quinn et al. 2021). Here, we use a marker-less CRISPR-HDR approach to introduce a defined resistance mutation into a susceptible genetic background.

As a proof-of-principle, we focus on the *rdl* (*‘resistance-to-dieldrin’*) locus. The A296S mutation is technically tractable, has a well-characterised molecular mechanism, and is associated with a strong resistance phenotype (Wondji et al. 2011). It also represents an informative case, having remained at high frequencies in African populations long after the withdrawal of dieldrin (Grau-Bové et al. 2020; Mugenzi et al. 2022). By introducing the A296S mutation into the susceptible *An. funestus* FANG strain, we quantify its contribution to resistance, examine its dominance across exposure regimes, and assess its effects on key life-history traits. By resolving both resistance effects and associated fitness trade-offs in a defined background, this provides insight into how such alleles may be maintained in natural populations, particularly where fitness costs are stage-specific or partially masked.

## RESULTS

### Marker-less CRISPR-HDR introduces A296S into *An. funestus*

To introduce the A296S mutation, we designed a CRISPR-HDR strategy utilising a guide RNA that recognises the wild-type allele but is unable to cut the edited *rdl* allele, as the causative mutation sits within the seed region. Provision of a plasmid-based homology template with the A296S mutation facilitates a repair event that prevents subsequent cutting by the Cas9-gRNA complex (Figure 1).

**Figure 1.**
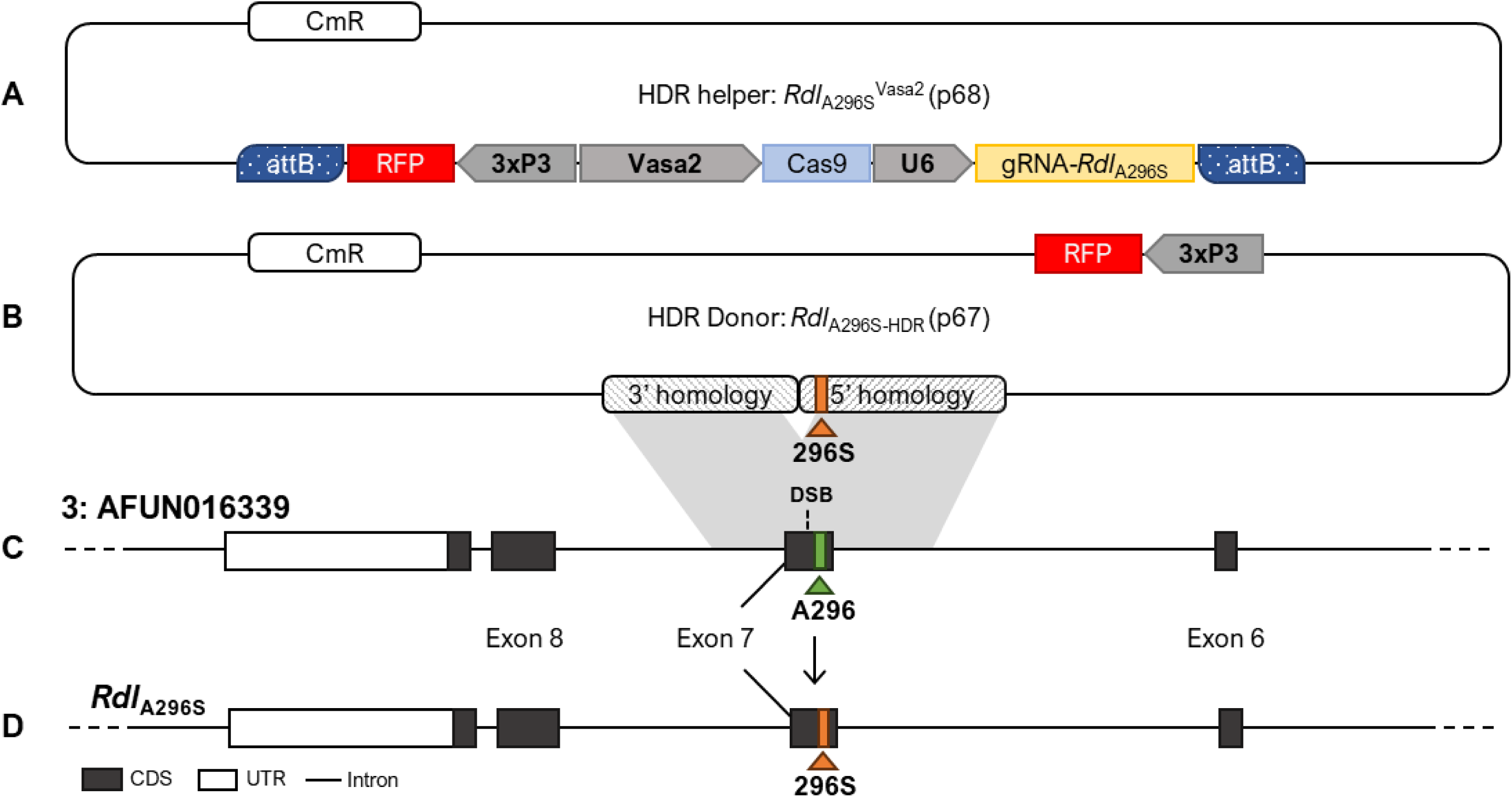
Schematic representation of the CRISPR-mediated HDR knock-in process for integration of the Rdl-A296S insecticide resistance allele into a susceptible *An. funestus* strain (FANG). DNA repair is mediated through the concurrent microinjection of two plasmid assemblies. A ‘helper’ plasmid (A) was designed to induce a double-stranded break (DSB) at the target locus (AFUN016339, C) upon expression, containing a source of Cas9 and gRNA, targeting the susceptible (296A) allele, under the control of the vasa2 and U6 promoters respectively. The ‘template’ plasmid (B) contained the Rdl-A296S SNP (296S) within the 5’ region of homology approximately 1kbp of the cut site. A second region of 3’ homology downstream of the cut site was included to facilitate the HDR process. Notably, a 3xP3::RFP marker cassette was included in the donor plasmid but outside the regions of homology, and so would not be integrated in a legitimate HDR event. Successful HDR events would be indicated by an alanine to serine substitution at the *Rdl* locus in the genome (D).

We employed a marker-less screening approach to avoid introducing additional genetic elements that could confound downstream phenotypic analyses. Because CRISPR-HDR events in the germline typically occur early in gametogenesis and thus result in ‘clusters’ of siblings that inherit the same edit (Quinn et al. 2021), positive founders were identified by screening a subset of progeny using TaqMan™ probes designed to distinguish the *rdl* allele from the wild-type, and the edit was subsequently recovered from non-sacrificed siblings (Supplementary Figure 1). Positive events were confirmed by Sanger sequencing. Two sub-lines (M25 and M33) were recovered from pooled crosses of transient males to wild-type females and established for propagation. As these crosses were performed *en masse*, we cannot exclude that both sub-lines derive from a single germline event (Supplementary Table 1). A third lineage (F4), derived from an independent female founder, was also recovered. Although F4 and M25 were eventually lost, M33 was successfully maintained for phenotypic analyses.

The recovery of multiple edited lineages from a modest number of injections demonstrates that marker-less HDR can be implemented efficiently in *An. funestus*, providing a practical framework for introducing defined resistance-associated alleles into a controlled genetic background for functional validation.

### Quantifying the effect size of A296S on dieldrin resistance

To assess the extent of resistance conferred by the A296S mutation, WHO tube insecticide bioassays were performed using a 1-hour exposure to the discriminating dose of 0.4% dieldrin on the M25 and M33 lines across genotypes (A/A, A/S, S/S). Assays were conducted with mixed and blinded genotypes using the offspring of heterozygote crosses with expected genotype frequencies of 1:2:1. Individual genotypes were determined post-exposure by qPCR. Mortality was analysed using a Bayesian generalised linear mixed model (GLMM) to accommodate inherent variability and complete separation; estimates reported below are model-predicted.

A total of 119 FANG-A/A, 251 FANG-A/S, and 70 FANG-S/S mosquitoes were tested across 22 replicates (tubes). Data were combined from two genome-edited sub-lines (M25 and M33) after confirming that the mutation behaved consistently across both backgrounds (Supplementary Figure 2). The observed departure from the expected 1:2:1 ratio – specifically the under-representation of homozygous resistant individuals – likely reflected a sampling bias towards early-emerging adults, which aligns with a significant developmental delay subsequently identified in FANG-S/S larvae (χ² = 19.65, df = 2, p < 0.0001).

Mortality differed strongly by genotype (Figure 2). Homozygous susceptible (FANG-A/A) individuals were fully susceptible (99.2% estimated mortality, 95% CrI: 97.5–100.0%), whereas homozygous resistant (FANG-S/S) individuals exhibited high levels of resistance (7.8% mortality, 95% CrI: 1.6–17.3%). Pairwise genotype contrasts confirmed the substantial protective effect of A296S on dieldrin resistance (Table 1). Heterozygotes (FANG-A/S) showed an intermediate phenotype (22.2% mortality, 95% CrI: 9.0–36.4%) (Figure 2), confirming that A296S confers strong resistance in the homozygous state with substantial protection in heterozygotes.

**Figure 2.**
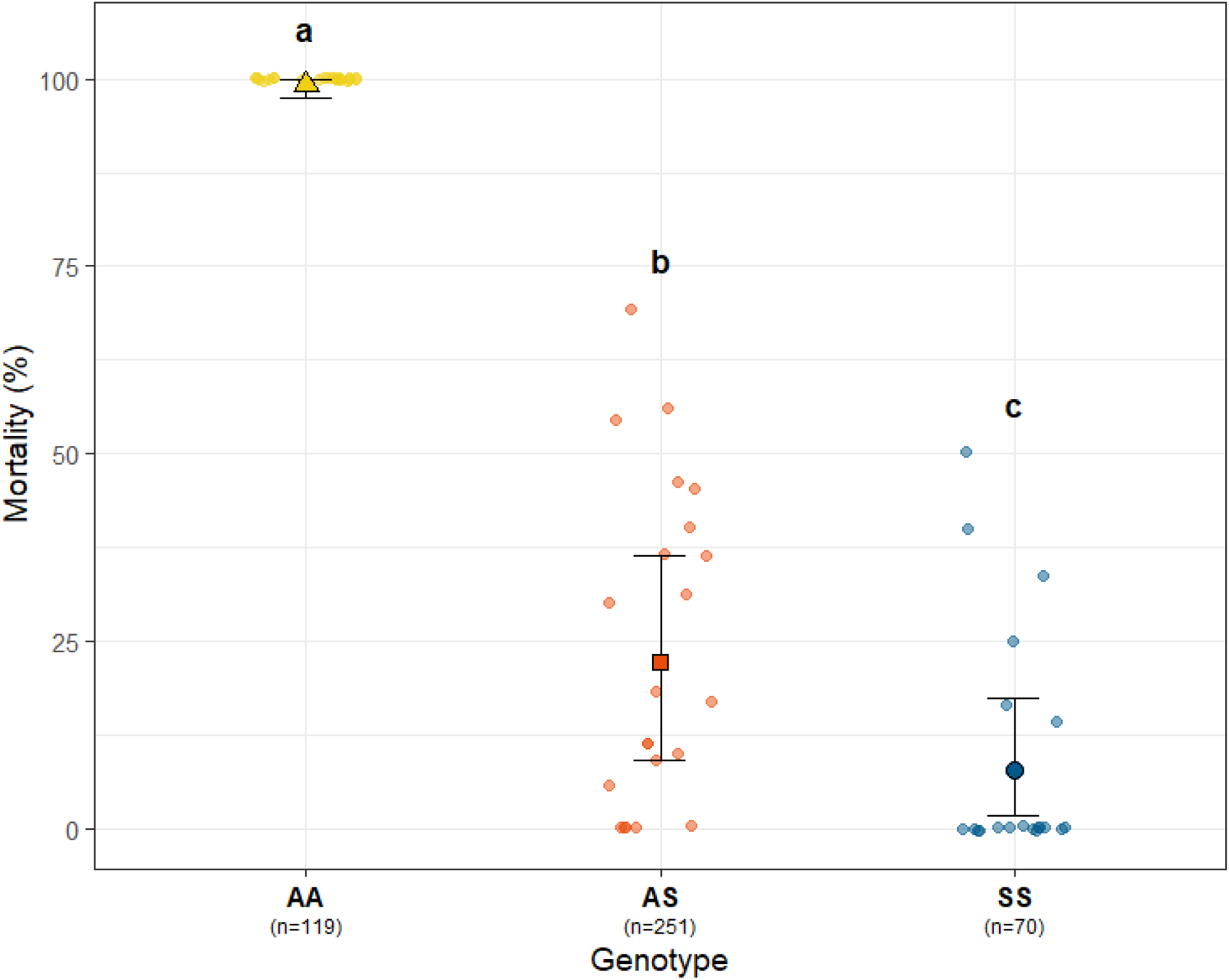
Mortality of adult female *Anopheles funestus* following exposure to 0.4% dieldrin for 1 h by *Rdl*-A296S genotype. Shapes and error bars indicate model-estimated posterior median mortality and 95% highest posterior density (HPD) intervals, averaged over sub-line, date, and tube. Background coloured points show observed tube-level mortality percentages; n numbers represent the total number of mosquitoes for that genotype. Genotypes with different letters indicate a statistically credible difference (95% CrI for the Odds Ratio does not overlap 1; Table 1). AA denotes homozygous susceptible (A296/A296), AS heterozygotes (A296/296S), and SS homozygous resistant (296S/296S) mosquitoes.

**Table 1.** Association between the Rdl-A296S mutation and mortality following exposure to insecticide at standard (1 hr) or reduced (30 minute) exposure times. Median Odds Ratios represent pairwise contrasts in mortality between genotype classes derived from Bayesian generalised linear mixed models (GLMMs), accounting for variation between assay dates and tubes. For the standard-dose assay, estimates were averaged across sub-lines. Values >1 indicate higher odds of mortality in the first-listed genotype relative to the second. Uncertainty is summarised using 95% highest posterior density (HPD) intervals; intervals including 1 indicate no clear difference in mortality odds between the contrasted genotypes. Relative toxic loads were approximated from the product of insecticide concentration and exposure duration, using 0.4% dieldrin for 1 hr as the reference. For the standard-dose dieldrin assay (0.4%, 1 hr), a prior sensitivity analysis using wider normal priors (SD = 5 and 10) confirmed that the genotype effect remained substantial, although the estimated odds ratios and their uncertainty increased considerably with weaker regularisation. A/A: homozygous susceptible; A/S: heterozygous; S/S: homozygous resistant; CrI: Credible Interval.

| *Insecticide* | *Exposure time* | *Toxic Load* | *Comparison* | *Median Odds Ratio* | *95% CrI (HPD)* |
|---|---|---|---|---|---|
| ***0.4% Dieldrin*** | ***1 hour*** | **1x** | **A/A vs. A/S** | 460.6 | 59.5 – 1753.6 |
|  |  |  | **A/A vs. S/S** | 1534.9 | 169.7 – 6760.5 |
|  |  |  | **A/S vs. S/S** | 3.33 | 1.17 – 7.13 |
| ***4% Dieldrin*** | ***1 hour*** | **10x** | **A/A vs. A/S** | 0.2 | 8.75 × 10<sup>−6</sup> – 3.21 |
|  |  |  | **A/A vs. S/S** | 43.2 | 0.22 – 763.4 |
|  |  |  | **A/S vs. S/S** | 216.9 | 0.28 – 10438 |
| ***4% Dieldrin*** | ***30 minutes*** | **5x** | **A/A vs. A/S** | 76.8 | 10.6 – 322 |
|  |  |  | **A/A vs. S/S** | 2852.3 | 161.6 – 16449 |
|  |  |  | **A/S vs. S/S** | 35.4 | 5.9 – 123 |
| ***4% DDT*** | ***30 minutes*** | **N/A** | **A/A vs. A/S** | 2.79 | 0.218 – 10.7 |
|  |  |  | **A/A vs. S/S** | 7.41 | 0.677 – 30.3 |
|  |  |  | **A/S vs. S/S** | 2.65 | 0.515 – 7.6 |

To evaluate resistance intensity, mosquitoes of all three genotypes were exposed to 4% dieldrin, representing 10x the standard WHO diagnostic dose (0.4%). We observed high mortality across all genotypes, with only 3 survivors out of 57 individuals (5.2%), all of which were homozygous resistant (Supplementary Figure 3). This precluded robust statistical analysis but is consistent with strong resistance in the homozygous state and little or no protection in heterozygotes under this extreme exposure. Reducing exposure time to 30 minutes improved discrimination between genotypes at this higher concentration (Figure 3, Table 1). FANG-A/A individuals remained highly susceptible (98.6% estimated mortality, 95% CrI: 93.6–99.8%), while FANG-S/S individuals retained strong resistance (2.4% mortality, 95% CrI: 0.0–9.1%). Heterozygotes showed increased mortality relative to the lower-dose assay (46.7% mortality, 95% CrI: 16.9–79.0%) (Figure 3), indicating reduced protection at higher toxic load.

**Figure 3.**
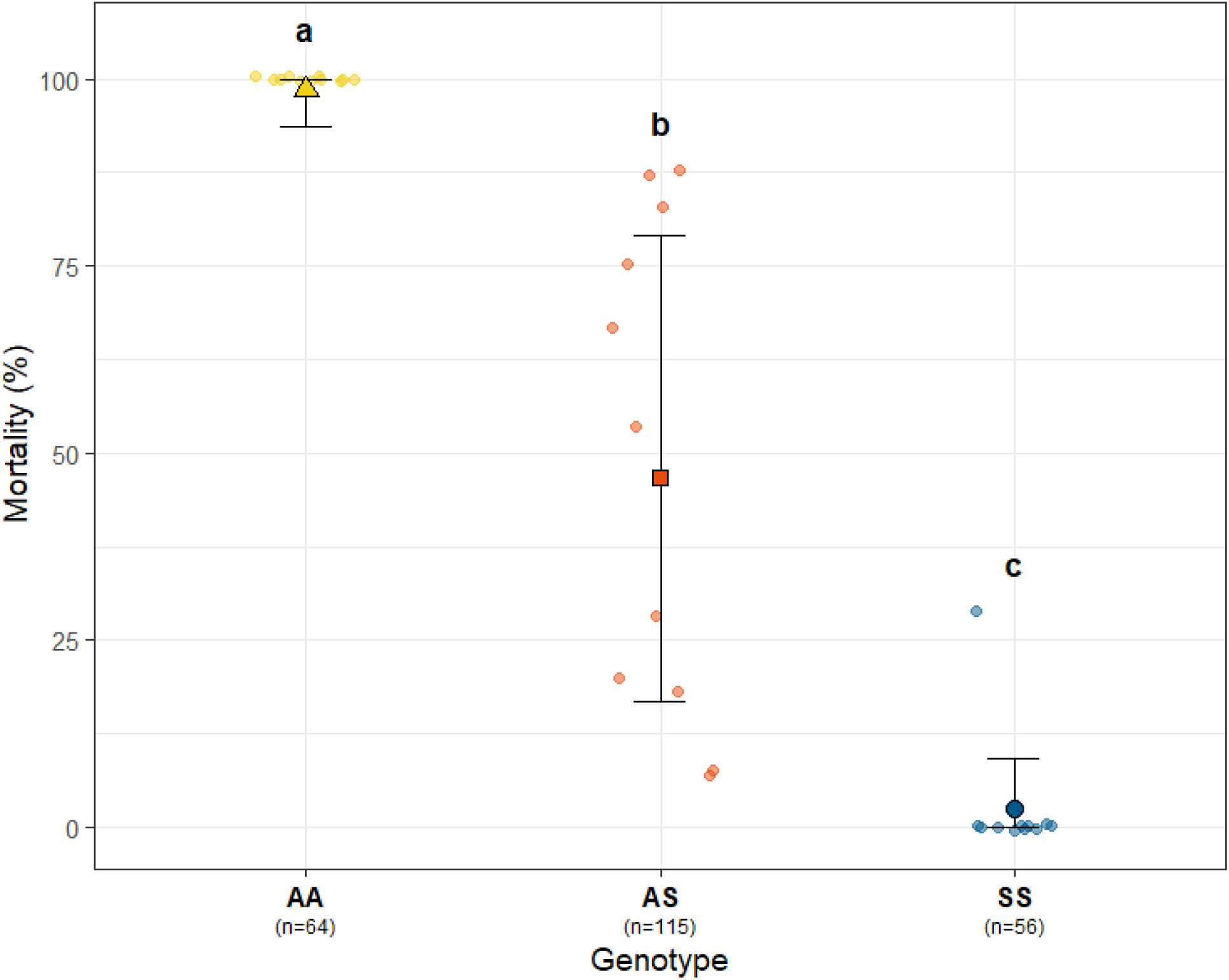
Mortality of adult female *Anopheles funestus* following exposure to 4% dieldrin for 30 min by *Rdl-A296S* genotype. Shapes and error bars indicate model-estimated posterior median mortality and 95% highest posterior density (HPD) intervals, averaged over sub-line, date, and tube. Background coloured points show observed tube-level mortality percentages; n numbers represent the total number of mosquitoes for that genotype. Genotypes with different letters indicate a statistically credible difference (95% CrI for the Odds Ratio does not overlap 1; Table 1). AA denotes homozygous susceptible (A296/A296), AS heterozygotes (A296/296S), and SS homozygous resistant (296S/296S) mosquitoes.

Consistent with this, the degree of dominance decreased across the exposure regimes tested, from D = 0.84 (95% CrI: 0.74–0.95) at 0.4% dieldrin to D = 0.55 (95% CrI: 0.42–0.68) at 4% dieldrin for 30 minutes (Table 2). This indicates that the protective effect of a single resistant allele is reduced under higher toxic load, with the phenotype shifting towards additivity within the range of exposure tested.

**Table 2.** Degree of dominance (D) of the Rdl-A296S resistance phenotype across insecticide treatments. Dominance coefficients were calculated on the relative fitness (survival) scale using the formula *D* = (*W_AS_* − *W_AA_*)⁄(*W_SS_* − *W_AA_*), where W represents the genotype-specific survival probability derived from the model-estimated (1 − *P_deat_*_ℎ_). On this scale, D = 0 represents a completely recessive trait, D = 0.5 an additive trait, and D = 1 a completely dominant trait. Values represent the posterior median and 95% Credible Intervals (CrI) generated from Bayesian generalised linear mixed models (GLMMs). All intervals were calculated using the Highest Posterior Density (HPD) method to ensure accurate coverage of the posterior distribution.

| ***Insecticide*** | ***Exposure time*** | ***Median D*** | ***95% CrI (HPD)*** |
|---|---|---|---|
| *0.4% Dieldrin* | *1 hour* | *0.84* | *0.74 – 0.95* |
| *4% Dieldrin* | *30 minutes* | *0.55* | *0.42 – 0.68* |

### Assessing Specificity and Cross-resistance

Given the historical use of both dieldrin and DDT in vector control and their structural similarities as organochlorines, we tested whether the Rdl-A296S mutation confers cross-resistance to DDT.

Exposure to 4% DDT for 1 hour was lethal across all genotypes (Supplementary Table 2). At a reduced exposure time of 30 minutes, a modest trend towards increased survival was observed in individuals carrying the 296S allele (Supplementary Figure 4); however, this effect was weak and inconsistent, with no credible separation between genotypes (Table 1).

Furthermore, no resistance was observed against permethrin or deltamethrin, widely used pyrethroids in long-lasting insecticidal nets, with complete susceptibility across all genotypes (Supplementary Table 2).

### Effect of A296S on general life history traits

A key consideration for any resistance mutation is its effect on fitness in the absence of insecticide selection, as this will influence its persistence following withdrawal of the selecting agent, with important consequences for resistance management through insecticide rotation. We therefore assessed the effects of Rdl-A296S on key life-history traits, including larval development and adult longevity.

### *Rdl-*A296S marginally increases larval development time in homozygosity

Larval development time was compared across genotypes to assess potential pleiotropic effects of the Rdl-A296S mutation.

Of 600 larvae, 514 (86%) survived to pupation. Genotypes were successfully determined for 482 individuals, which did not deviate from expected Mendelian ratios (χ² = 3.99, df = 2, p = 0.136), providing no evidence of genotype-dependent differences in survival to pupation.

Overall larval development time was tightly distributed (median 12 days, IQR: 11–12; Supplementary Table 3). Males developed slightly faster than females (median 11 vs 12 days), but the difference was not statistically significant (hazard ratio [HR]: 1.12, p = 0.233; Supplementary Table 4). Susceptible (FANG-A/A) and heterozygous (FANG-A/S) larvae showed similar developmental profiles (median 11 days), whereas homozygous resistant (FANG-S/S) larvae exhibited a shift towards later pupation (median 12 days, IQR: 11–13), with an increased proportion of individuals pupating at days 14–15 relative to the susceptible reference (Figure 4, Supplementary Table 3).

**Figure 4.**
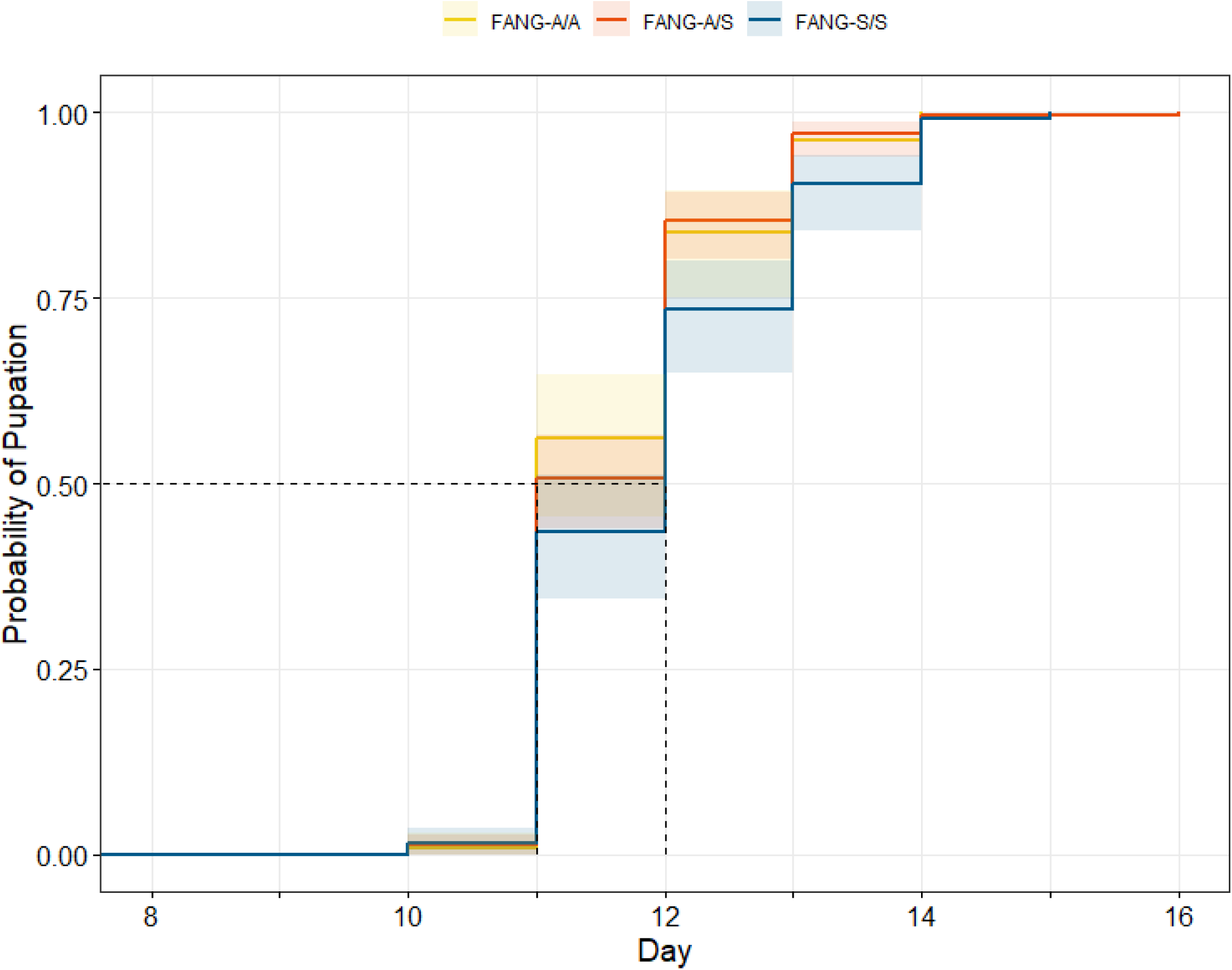
Cumulative pupation rate of Anopheles *funestus* genotypes estimated through Kaplan-Meier survivorship, showing the time-to-pupation (days post-hatching) for the *Rdl*-A296S mutation on the FANG genetic background. Curves represent the cumulative proportion of larvae reaching the pupal stage for homozygous susceptible (FANG-A/A; yellow), heterozygous (FANG-A/S; red), and homozygous resistant (FANG-S/S; blue) individuals. Data are derived from 482 individuals with confirmed genotypes (nA/A = 105, nA/S = 241, nS/S = 136). Shaded areas represent 95% confidence intervals.

A Cox proportional hazards model confirmed that this delay was restricted to the homozygous state (HR = 0.74, 95% CI: 0.57–0.95, p = 0.021), with no difference between heterozygotes and susceptible individuals (Supplementary Table 4). As the event modelled was pupation, a hazard ratio (HR) below 1 indicates a reduced rate of pupation and therefore slower development relative to the reference genotype (FANG-A/A). The observed genotype frequencies did not deviate significantly from the expected Mendelian ratio (*χ*^2^ = 3.99, df = 2, p = 0.136), providing no evidence of genotype-dependent differences in survival to pupation.

### Effect of Rdl-A296S on Adult longevity

Adult longevity was assessed in single-sex, mixed-genotype cages (n = 878), using offspring derived from intercrossed heterozygotes (Supplementary Table 5). At the start of the assay, genotype frequencies followed the expected 1:2:1 Mendelian ratio (χ² = 0.039, df = 2, p = 0.981), indicating that the *Rdl*-A296S mutation did not incur significant genotype-dependent mortality during larval or pupal development. Survival was then tracked daily with genotypes determined post-mortem.

Adult survival was strongly influenced by sex, with males surviving considerably longer than females under assay conditions (median 57 vs 39 days; Figure 5, Supplementary Table 5). Importantly, there was no evidence of a longevity cost associated with the 296S allele. Rather, a consistent trend towards increased longevity in heterozygous and homozygous resistant individuals relative to susceptible individuals was evident from survivorship curves and summary statistics (Figure 5, Supplementary Table 5). Cox proportional hazards supported this observation, with heterozygotes exhibiting a modest survival advantage (HR = 0.83, 95% CI: 0.70–0.97, p = 0.023), while homozygous resistant individuals showed a similar but non-significant trend (HR = 0.87, p = 0.163; Supplementary Table 6).

**Figure 5.**
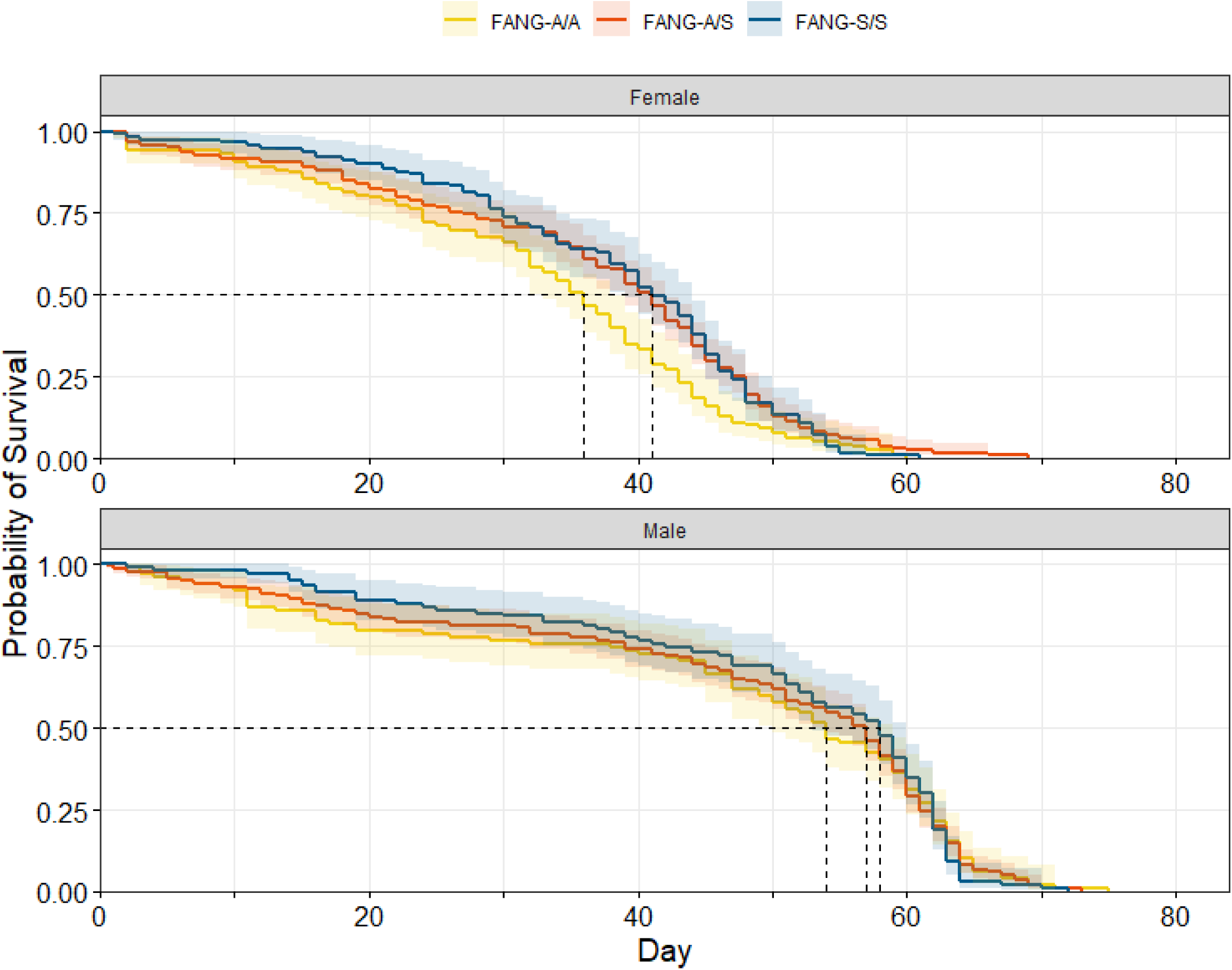
Adult survivorship of *Anopheles funestus* genotypes containing the Rdl-*A296S* mutation, estimated through Kaplan-Meier survival curves for susceptible (FANG-A/A; yellow), heterozygous (FANG-A/S; red) and homozygous resistant (FANG-S/S; blue) mosquitoes, stratified by sex. Plots illustrate the probability of survival over time (days post-emergence) for females (top) and males (bottom). Shaded areas represent 95% confidence intervals. Median survival significantly differed between sexes at 57 days for males and 39 days for females (p < 0.001). Overall, adult heterozygous (FANG-A/S) individuals exhibited a significant survival advantage over the susceptible (FANG-A/A) reference (HR: 0.83, p = 0.023). Dashed lines indicate the median survival time (0.5 probability) for each genotype within each sex. Data are aggregated from two independent replicates (n = 878).

## DISCUSSION

This study provides the first in vivo functional validation of an insecticide resistance mutation in *Anopheles funestus*, a major malaria vector that has historically proven recalcitrant to germline transformation. By engineering a marker-less strain on a fully susceptible genetic background, we have decoupled the *Rdl* locus from the confounding metabolic and structural variation present in wild populations. This provides clear evidence that the A296S substitution alone is sufficient to confer high-level phenotypic resistance to dieldrin.

Beyond establishing causality, our results define the phenotypic properties of this mutation under controlled conditions. Resistance is strong but context-dependent: at lower exposures, survival is high in both homozygous resistant and heterozygous individuals, whereas at higher exposures resistance remains robust in homozygotes but heterozygote protection is reduced, shifting the phenotype towards additivity. This demonstrates that the degree of dominance varies across the exposure regimes tested, although the individual contributions of insecticide concentration, exposure duration, and sub-line composition cannot be resolved. In the field, where insecticide residues are likely to be non-uniform or decaying over time, even this partial protection may confer a substantial selective advantage on heterozygotes under some exposure conditions (South et al. 2020).

Consistent with this, the persistence of *Rdl* in African vector populations for decades following the withdrawal of dieldrin has long been noted, and a range of explanations have been proposed, including environmental persistence of dieldrin, continued or illicit use, cross-resistance to other compounds, or minimal fitness costs such that there is little selective pressure to reduce frequency of the allele (Grau-Bové et al. 2020; Mugenzi et al. 2022). We find no evidence for operationally relevant cross-resistance to DDT or to widely used pyrethroids, indicating that the effect of A296S is highly target-specific. At the same time, the mutation imposes a modest, recessive cost during larval development, with no detectable reduction in adult survival and, at most, a small and variable heterozygote advantage. Taken together, these results suggest that the net fitness effect of A296S is close to neutral under laboratory conditions, particularly in the heterozygous state. Our findings therefore argue against cross-resistance to DDT or pyrethroids as a mechanism for maintaining this allele and are consistent with a scenario in which the direct fitness costs of A296S are limited, recessive, and partially offset across life stages, reducing the strength of purifying selection once insecticide pressure is removed. While this does not resolve the relative contributions of these mechanisms in the field, it provides a necessary baseline for evaluating them.

It is important to note that these assays capture only a subset of potential fitness effects. As a component of GABAergic signalling, *Rdl* may influence a broader range of traits, including behaviour and mating interactions (Platt et al. 2015; Rowland 1991a, 1991b), which are not captured in laboratory assays of development and survival. More generally, fitness costs are often difficult to infer from laboratory studies alone, particularly when field-realistic contexts are not reproduced (ffrench-Constant and Bass 2017). In our study, the use of a sugar-only diet and sex-separate cages removed key aspects of normal ecology, including blood-feeding, mating, and reproduction, all of which may impose sex-specific energetic costs (Dao et al. 2010). As a result, the observed survival patterns may not fully reflect longevity under standard colony maintenance or natural settings. A comprehensive assessment of fitness consequences would therefore require evaluation under more complex and ecologically relevant conditions, for example through semi-field studies tracking allele dynamics over time. Recent evidence from the *An. gambiae* species complex suggests that *Rdl* resistance mutations can enhance auditory function and mating-related behaviours (Sangbakembi-Ngounou et al. 2026). If these effects are conserved in *An. funestus*, they could confer fitness advantages in particular ecological contexts, potentially contributing to the persistence of resistance alleles long after the withdrawal of dieldrin.

While field studies associate the A296S mutation with very strong survival at high-intensity exposures, our results show that the mutation provides much weaker protection in isolation. In a population of field-derived *An. funestus* from Nigeria exposed to 4% dieldrin for 1 hour, post-bioassay genotyping showed that survivors were overwhelmingly homozygous resistant, with a smaller contribution from heterozygotes, whereas mortality was concentrated among homozygous susceptible mosquitoes, with few heterozygotes and no homozygous resistant individuals (Djouaka et al. 2016). By contrast, under the same exposure regime, our edited line showed no survival in heterozygotes and only limited protection in homozygous resistant mosquitoes. This marked discrepancy indicates that the apparent effect size of the allele in wild populations may be amplified by genomic background. In particular, the lower resistance levels observed in our edited line relative to some field estimates are consistent with the presence of additional co-occurring resistance mechanisms in natural populations. In *An. funestus*, the restoration of susceptibility by PBO in field populations, together with resistance to multiple insecticide classes in the same population (Djouaka et al. 2016), and the documented upregulation of detoxification genes in the same region (Atoyebi et al. 2020), supports the view that metabolic and other background resistance mechanisms can substantially modify the observed phenotype associated with *Rdl*. By isolating A296S on a defined susceptible background, our study provides a quantitative baseline for the resistance effect of the mutation in isolation, separated from the confounding influence of co-occurring mechanisms.

These findings also have implications for molecular surveillance. While genetic markers are often used as proxies for resistance in field populations, their utility depends on knowing the precise effect size of individual mutations (Donnelly, Isaacs, and Weetman 2016). By isolating A296S on a defined genetic background, this study provides a quantitative benchmark for resistance intensity and demonstrates the importance of functional validation in linking genotype to phenotype (Grigoraki et al. 2021; Williams et al. 2022). For *An. funestus*, the marker-less editing framework presented here is well suited to future studies of interactions between resistance-associated variants, including markers of metabolically mediated resistance such as *GSTe2*-L119F (Riveron et al. 2014). The same platform also provides a route to test the recently described L976F (L995F in *An. gambiae*) allele (Odero et al. 2024), to elucidate why this canonical *kdr* mutation appeared to be absent in *An. funestus* for so long despite intense pyrethroid selection (Irving and Wondji 2017; Menze et al. 2016).

This work establishes a practical framework for precise genome editing in *An. funestus*, enabling the marker-less introduction of defined, field-relevant alleles via homology-directed repair and overcoming a long-standing barrier to functional genetics in this species. Resistance phenotypes emerge from interactions between causal mutations, genetic background, insecticide exposure, and ecological context. Even mutations with large effect sizes do not act in isolation, and their phenotypic expression will depend on dose, delivery, behaviour, and interactions with other loci. More broadly, functional validation in controlled backgrounds, as demonstrated here, provides a foundation for disentangling these effects and for linking genotype to phenotype in a way that is relevant to operational decision-making by underpinning resistance monitoring and management.

## ACKNOWLEDGEMENTS

The authors are grateful to Amy Guy and colleagues at the Liverpool Insect Testing Establishment (LITE) for their expertise in mosquito rearing and colony maintenance. We also thank Zachary Stavrou-Dowd, Jasmine Morgan, and Poppy Pescod for their technical assistance and helpful discussions, Jessica Carson for her laboratory assistance, Abdullahi Muhammad and Helen Irving for their training and guidance, and Frank Mechan for his statistical advice.

This work was supported by funding from the Medical Research Council (MRC).

## ETHICS STATEMENT

The use of human blood in our mosquito feeding protocols was conducted under the authority of License Number 12548, granted to the Liverpool School of Tropical Medicine (LSTM), Pembroke Place, Liverpool, L3 5QA. This license is issued under Section 16(2)(e)(ii) of the Human Tissue Act 2004 and held by Professor Jonathan Ball.

All work with genetically modified mosquitoes was carried out in containment level 2 insectaries under GMM 113 and GMM 141 in accordance with UK legislation on Genetically Modified Organisms 2005 and 2014 under Dr Gareth Lycett and Prof Tony Nolan.

## METHODS

### Mosquito strains and rearing

Mosquitoes (*Anopheles funestus*, FANG strain) (Hunt et al. 2005) were maintained under standard insectary conditions (26°C ± 2°C, 70% ± 10% RH, 12:12 h L:D). Larvae were reared in purified water at a density of 0.24 larvae/mL and fed ground fish food (TetraMin Tropical Flakes, Tetra, Blacksburg, VA, USA) at increasing daily dosages (100–233 µg/larva). Adults were provided with 10% sucrose solution *ad libitum.* Colony maintenance was performed by twice-feeding adult females using a Hemotek Membrane Feeding System (Hemotek Ltd., Blackburn, UK) with screened human blood. For the establishment of nascent gene edited lines (G0 – G3), small groups of females were fed on a human volunteer, in accordance with LSTM’s code of practice for arm-feeding *Anopheles* mosquitoes, or a modified Hemotek system using small, customised mesh-covered chambers to maximise probing efficiency at low mosquito densities.

### CRISPR/Cas9 plasmid design and construction

CRISPR/Cas9 components were designed using the *An. funestus* AFunF3_3v52 reference genome (Neafsey et al. 2015). Candidate 20-bp sgRNA sites targeting exon 7 of the GABA receptor gene (AFUN016339) were identified using CHOPCHOP v3 (Labun et al. 2019). The helper plasmid (p68, Figure 1A) was derived from the vasa-Cas9 expression vector p165 (KU189142) (Hammond et al. 2016), which expresses a human-codon-optimised SpCas9 under the *An. gambiae vasa2* promoter (Papathanos et al. 2009) and carries a U6-driven sgRNA cassette (Hwang et al. 2013). An *Rdl*-specific spacer was generated by annealing oligos TN76/TN77 (Supplementary Table 7) and cloned into p165 via *BsaI* Golden Gate cloning.

The homology-directed repair (HDR) donor plasmid (p67, Figure 1B) was constructed via Gibson Assembly (New England Biolabs). Regions of homology (∼1 kb) flanking the cut site were amplified from FANG gDNA using primers TN72–TN75 (Supplementary Table 7). The A296S substitution (GCA→TCA) was introduced through overlapping primers TN73/TN74 (Supplementary Table 7). The donor was designed with a 3xP3-RFP marker cassette in the p165 backbone, located outside the homology arms and was therefore not expected to integrate during bona fide HDR events, allowing for the identification of transients and illegitimate integration, and subsequent identification of marker-less HDR events.

### Embryo microinjection and line establishment

Embryo microinjections were performed on the susceptible FANG strain following established protocols (Benedict 2007; Fuchs, Nolan, and Crisanti 2012; Quinn et al. 2021). Briefly, gravid females were induced to oviposit in darkness, and freshly laid embryos were aligned on nitrocellulose paper for injection at the posterior pole. The injection mix contained 300 ng/µL donor and 150 ng/µL helper plasmids in 1x injection buffer (0.1 mM sodium phosphate pH 6.8; 10 mM KCl). Hatched G0 larvae were screened for transient RFP expression within 48 hours of hatching, and surviving transient adults were outcrossed to the wild-type FANG strain in sex-specific groups.

G1 progeny from individual females (isofemale families) were pre-screened for the A296S allele by TaqMan genotyping as pooled larvae; where positive families were detected, remaining siblings were individually genotyped non-destructively using pupal cases (Supplementary Figure 1). Three edited lineages (M25, M33 and F4) were recovered and propagated through successive outcrossing and intercrossing. M25 and M33 were derived from pooled crosses involving transient G0 males, whereas F4 originated from an independent transient G0 female. Both M25 and M33 contributed to the initial 0.4% dieldrin bioassays; however, M25 was eventually lost following a colony collapse at approximately G10–G11. F4 exhibited poor colony viability over several generations and was similarly lost around G10–G11 without contributing to phenotypic analyses. Subsequent assays were therefore conducted using sub-line M33 only.

### Genotyping and molecular validation

Genomic DNA for screening and genotyping was extracted from larvae, pupal cases, or adults using a rapid STE-based extraction. Briefly, samples were homogenised in STE buffer (0.1 M NaCl, 10 mM Tris-HCl, 1 mM EDTA, pH 8.0) (12–40 µL depending on sample type) followed by incubation at 95°C for 60 minutes. Marker-less genotyping for the A296S substitution was performed using a Custom TaqMan SNP Genotyping Assay (Applied Biosystems) using allele-specific probes (Supplementary Table 8). Each 12 µL reaction contained 1x SensiMix II Probe Mix (Bioline), 1x TaqMan assay mix (Applied Biosystems) and 2.4 µL of gDNA. Thermal cycling was performed on an AriaMx Real-Time PCR System (Model no. G8830A, Agilent Technologies) under the following conditions: 95°C for 10 min, 40 cycles of 95°C for 10s and 60°C for 45s, with allelic discrimination performed using standard FAM (resistant 296S) and VIC (susceptible A296) probes. The presence of the A296S (GCA→TCA) substitution was confirmed by Sanger sequencing (GeneWiz, UK) following full gDNA extraction (Wizard® Genomic DNA extraction kit, Promega).

### Insecticide susceptibility phenotyping

Phenotypic resistance to insecticides was evaluated using standard WHO tube bioassays (World Health Organization 2016) on non-blood-fed adult females (2-5 days old). To assess the contribution of the A296S mutation to resistance intensity and dominance plasticity, FANG-A296S mosquitoes were exposed to dieldrin at two concentrations: the standard discriminating dose (0.4%) for 60 minutes and a high-intensity dose (4%) for 30 or 60 minutes. Cross-resistance was similarly evaluated using 4% DDT, 0.75% permethrin, and 0.05% deltamethrin. In all assays, up to 25 mosquitoes were exposed per tube to insecticide-treated papers (Universiti Sains Malaysia, or prepared in-house following LITE SOPs). Because adult females were limiting in some experiments, untreated, age-matched males were run in parallel as sentinel tubes to monitor for aberrant insectary conditions and handling-associated mortality. These tubes were used only as procedural quality-control checks and not to estimate baseline mortality in the insecticide-exposed female cohorts. Mortality was scored 24 hours post-exposure. Assays were performed on the unscreened offspring of heterozygotes; all individuals (dead and alive) were subsequently genotyped post-assay using TaqMan qPCR to determine genotype-specific mortality without prior selection bias.

### Fitness assays

#### Larval development

Life-history traits (larval development and adult longevity) were compared across *Rdl* genotypes to identify potential fitness trade-offs. For larval development, eggs were collected from heterozygous females following two blood meals, hatched, and larvae were distributed into three trays at a density of 200 larvae per tray (n = 3 trays). Larvae were then reared *en masse* under standard insectary conditions until pupation. Pupation was scored daily; pupae were first sexed and pupal cases were retained for TaqMan genotyping to assign genotype (FANG-A/A, A/S, and S/S) to each individual’s time to pupation.

#### Adult longevity

For adult longevity, larvae were first reared under standard insectary conditions as described above. To assess intrinsic longevity and eliminate mating-associated mortality, pupae were sexed and adults were housed in single-sex cages (up to 50 mosquitoes per cage) grouped by day of pupation. Adults were maintained without blood feeding and provided 10% sucrose *ad libitum*. Mortality was recorded daily until all adults had died, and dead individuals were genotyped post-mortem to assign survival outcomes to *Rdl* genotype. Adults that failed to emerge or were lost during handling were excluded from the analysis. Adult emergence was defined as pupation day +2 days, based on established life-history parameters for this strain (Zengenene et al. 2021). Two independent replicate experiments were conducted on different dates.

### Statistical analyses

#### Bioassay Modelling Framework

Statistical analyses were performed in R v4.3.1. Insecticide bioassay data were analysed within a Bayesian framework to accommodate inherent variability and potential complete separation arising from outcomes such as 100% mortality. Genotype-specific mortality was modelled using Bayesian generalised linear mixed models implemented in the *brms* package v2.23.0 (Bürkner 2017).

For all bioassays, the response variable was specified as the number of dead mosquitoes out of the total tested per tube and modelled using a binomial distribution with a logit link. The baseline model for the standard discriminating dose (0.4% dieldrin, 1 hr) included genotype and sub-line (M25 or M33) as fixed effects. Random intercepts were included for exposure tube and experimental date. Each tube was treated as an independent experimental unit and assigned a unique TubeID across the full study period, ensuring that tubes used on different dates were modelled as distinct observations rather than as repeated measures. Date was included separately to account for day-to-day variation in insectary conditions.

Models were estimated using four Markov chain Monte Carlo (MCMC) chains, each run for 4,000 iterations with a 2,000-iteration warm-up period, yielding a total of 8,000 post-warm-up samples. To improve sampling efficiency and facilitate convergence in datasets exhibiting complete separation, such as 100% mortality in FANG-A/A mosquitoes, the `adapt_delta` parameter was set to 0.95 and increased to 0.999 for the 4% dieldrin, 1 hr bioassay. Weakly informative priors were applied throughout: normal(0, 2.5) priors for the intercept and fixed effects, and a Cauchy(0, 2) prior for the standard deviations of random effects.

#### Model Comparison and Line Pooling

To determine how the two sub-lines should be handled in the 0.4% dieldrin, 1 hr bioassay, models were compared using leave-one-out cross-validation (LOO-CV) implemented in the *loo* package v2.8.0 (Vehtari, Gelman, and Gabry 2017). An additive model containing genotype and sub-line as main effects was compared with a model including a genotype-by-sub-line interaction. This tested whether the phenotypic effect of A296S differed between the two sub-lines. Following LOO implementation, the models showed equivalent predictive performance, with an expected log predictive density (ELPD) difference of <2. The additive model was therefore selected on the basis of parsimony, supporting the pooling of data across sub-lines while retaining sub-line as a fixed effect.

Whereas most models were fitted using 4,000 iterations per chain and an `adapt_delta` value of 0.95, the model for the 4% dieldrin, 1 h bioassay required more stringent sampling parameters to achieve convergence. Complete mortality among both susceptible FANG-A/A and heterozygous FANG-A/S mosquitoes produced a sharply defined posterior and sampling difficulties. The final model was therefore fitted using 10,000 iterations per chain, including 5,000 warm-up iterations, with `adapt_delta` increased to 0.999 and `max_treedepth` set to 15. These settings removed divergent transitions and improved Hamiltonian Monte Carlo sampling.

#### Posterior Inference and Dominance Estimation

To assess sensitivity to prior specification, the standard-dose dieldrin model was additionally fitted using wider normal priors (SD = 5 and 10) for the fixed-effect coefficients, compared with the original prior (SD = 2.5), while retaining the original priors for the intercept and random effects.

Genotype-specific mortality probabilities and pairwise Odds Ratios (OR) were derived from the posterior distributions using the *emmeans* package v2.0.0 (Lenth and Piaskowski 2025). Uncertainty was summarised using 95% Highest Posterior Density (HPD) intervals. The degree of dominance, (*D*), was calculated on the survival scale (*W* = 1 − *P_dead_*) following Stone (1968):

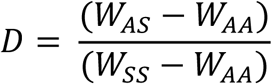

where *D* = 0 represents a completely recessive trait, *D* = 0.5 an additive trait and *D* = 1 a completely dominant trait. Genotype-specific survival probabilities (*W*) were derived from model-estimated mortalities.

### Life-history Analysis

Larval development time was analysed as a time-to-event outcome, with pupation defined as the event, using Cox proportional hazards regression implemented in the *survival* package v3.8-3 (Therneau 2024b) in R. The analysis was restricted to 482 individuals with successfully confirmed genotypes. To quantify the effect of the A296S mutation on developmental rate, we fitted a Cox proportional hazards model using the `coxph()` function, with genotype (FANG-A/A, FANG-A/S and FANG-S/S) and sex (female or male) included as fixed effects. Homozygous susceptible FANG-A/A mosquitoes and females were specified as the reference groups.

Differences in developmental rate were summarised as hazard ratios (HRs), with an HR <1 indicating a lower instantaneous probability of pupation at any given time point and therefore slower development relative to the reference group. The proportional hazards assumption was assessed using scaled Schoenfeld residuals with the `cox.zph()` function, with no evidence of violation detected (global test: χ² = 0.73, df = 3, p = 0.87). Cumulative probability of pupation over time was visualised using Kaplan–Meier estimates transformed as 1 − *S*(*t*), generated with the *survminer* package v0.5.1 (Kassambara, Kosinski, and Biecek 2025).

Adult survival data from two independent replicates were combined (n = 878) and analysed using a mixed-effects Cox proportional hazards model implemented in the *coxme* package v2.2-22 (Therneau 2024a). Sex and genotype were included as fixed effects, with females and FANG-A/A specified as the respective reference groups, and replicate included as a random intercept to account for between-replicate variation. An exploratory model including a sex-by-genotype interaction was also evaluated, but the additive model was retained for primary reporting based on its lower Akaike information criterion (AIC = 235.9 versus 237.8). Median survival times and 95% confidence intervals were derived from Kaplan-Meier estimates, with survival curves generated using *ggsurvplot()* and faceted by sex and grouped by genotype. The proportional hazards assumption was assessed using Schoenfeld residuals from a supplementary cox model stratified by replicate, revealing evidence of non-proportionality for sex and genotype (global test: χ² = 19.25, df = 3, p < 0.001). Hazard ratios from the primary mixed-effects model were therefore interpreted cautiously as overall summaries of survival differences rather than constant effects throughout the observation period.

## SUPPLEMENTARY MATERIAL

**Supplementary Figure 1.**
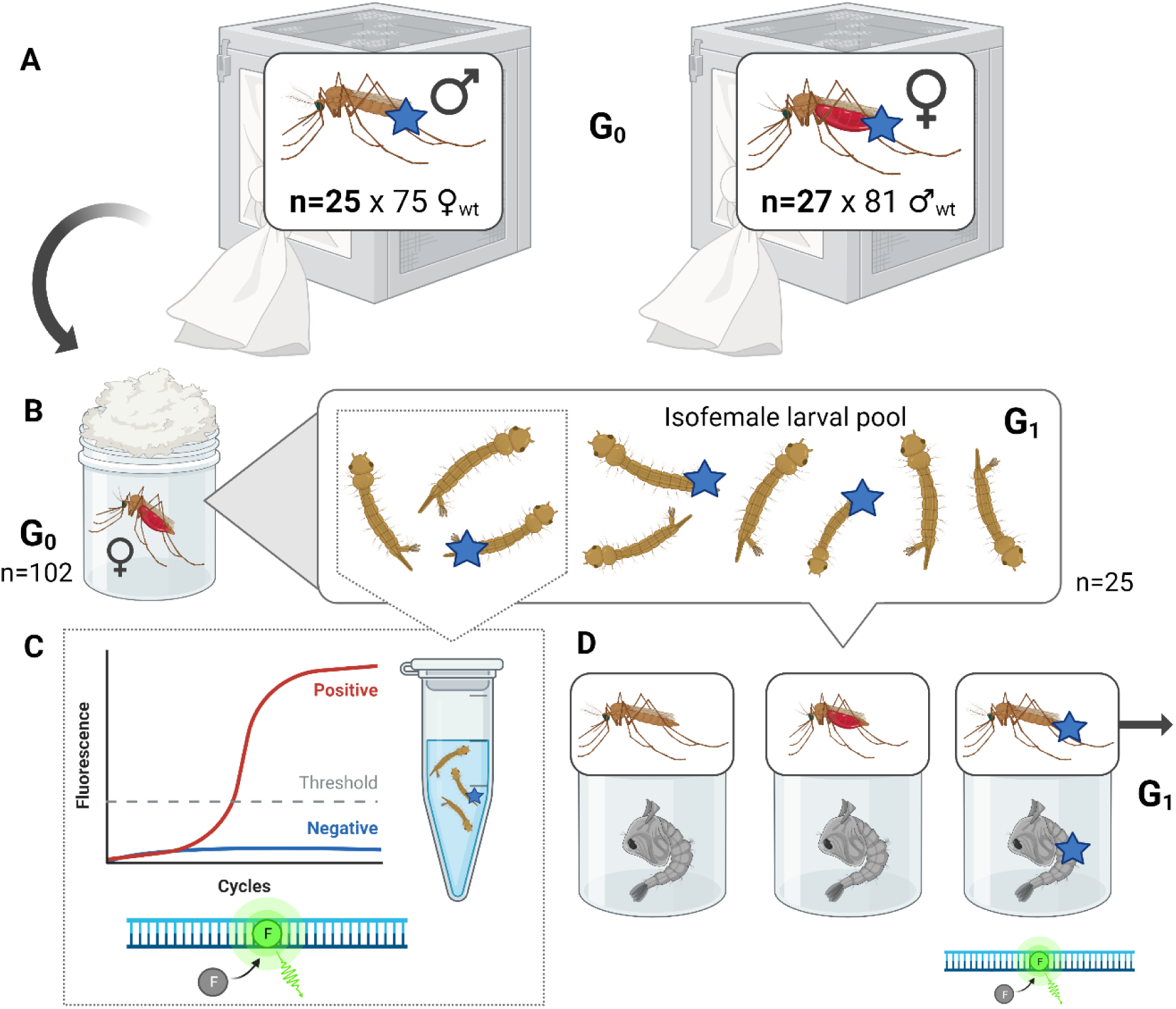
TaqMan™ qPCR-based screening strategy for recovering markerless edited FANG-A296S mosquitoes. Following microinjections of embryos, surviving adults exhibiting transient RFP expression were crossed with wild-type, and females isolated to oviposit. A subset of the progeny was screened for the mutation using TaqMan qPCR, and if the mutation was identified, the pupal cases of all the remaining siblings in the isofemale pool were screened and then crossed again with wildtype to establish the line.

**Supplementary Figure 2.**
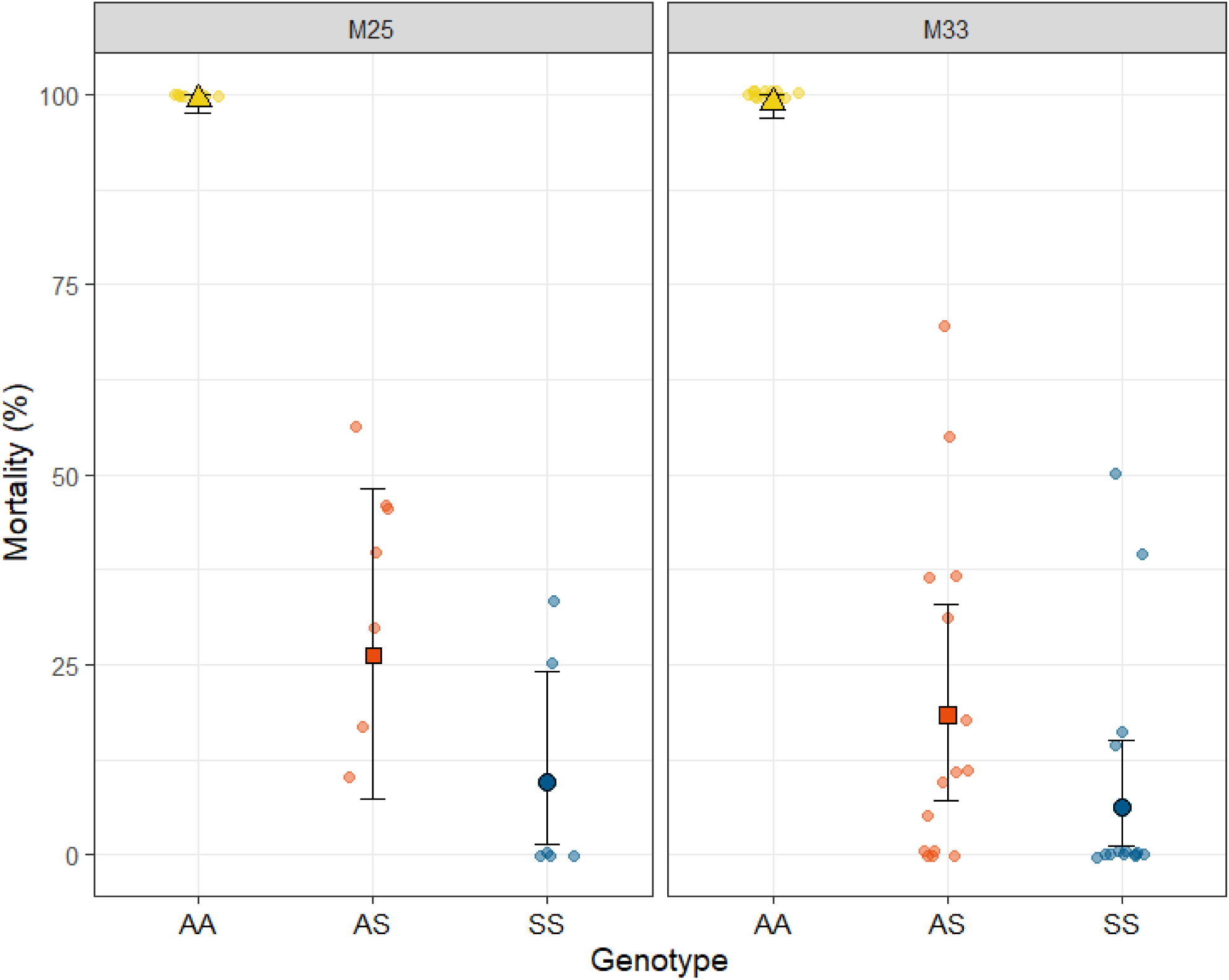
Mortality of adult female *Anopheles funestus* following exposure to 0.4% dieldrin for 1 hr by *Rdl*-A296S genotype, stratified by line (“M25” and “M33”). Shapes and error bars indicate model-estimated posterior median mortality and 95% highest posterior density (HPD) intervals, averaged over date and tube. Background coloured points show observed tube-level mortality percentages; n numbers represent the total number of mosquitoes for that genotype. AA denotes homozygous susceptible (A296/A296), AS heterozygotes (A296/296S), and SS homozygous resistant (296S/296S) mosquitoes. Statistical comparison using Leave-One-Out Cross-Validation (LOO-CV) confirmed that the phenotypic effect of the A296S mutation was consistent across both sub-lines, justifying the use of a pooled additive model for primary reporting. Sample sizes for each group are as follows: Line M25 (nAA = 40, nAS = 76, nSS = 21); Line M33 (nAA = 79, nAS = 175, nSS = 49).

**Supplementary Figure 3.**
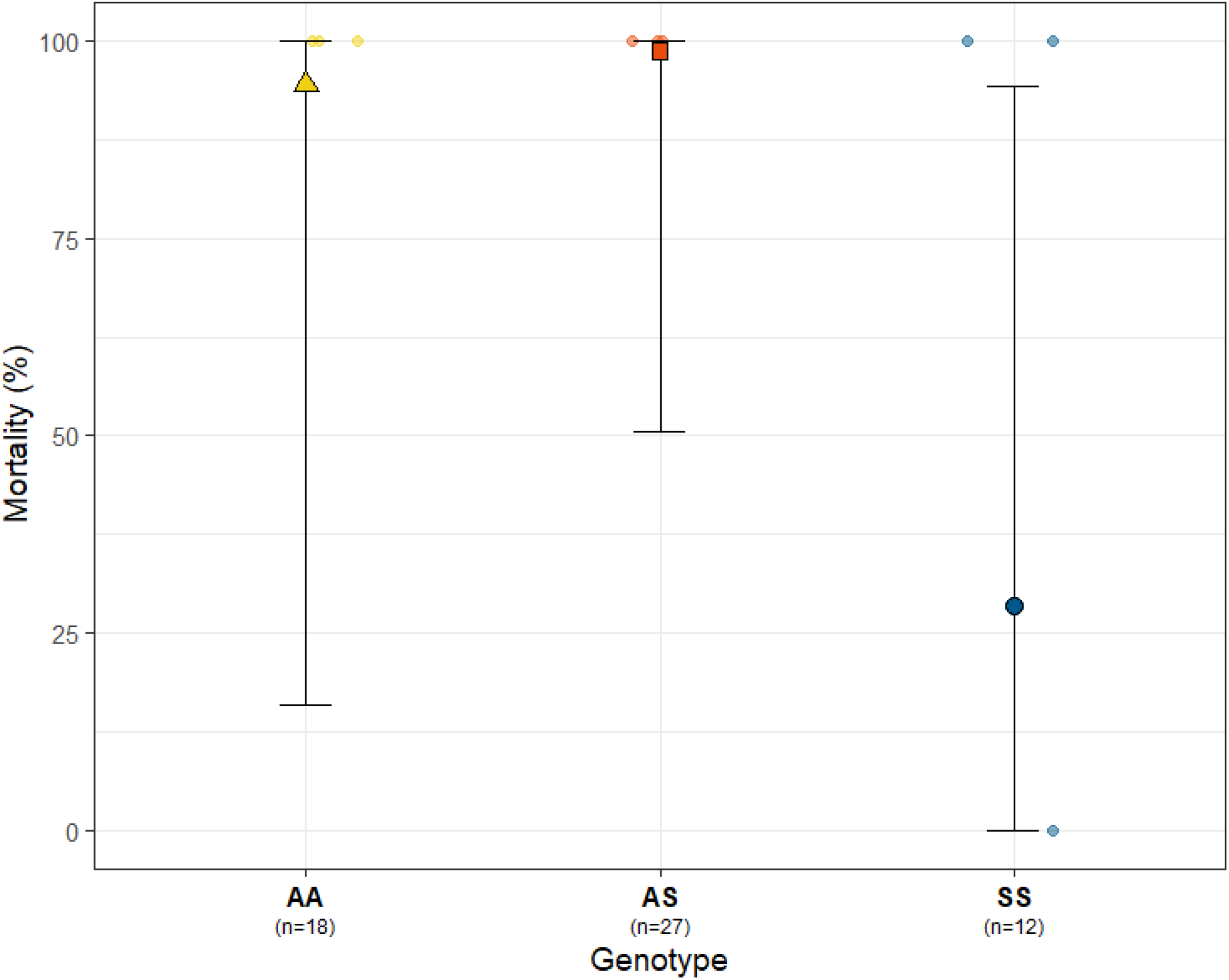
Mortality of adult female *Anopheles funestus* following exposure to 4% dieldrin for 1 hr by *Rdl*-A296S genotype. Shapes and error bars indicate model-estimated posterior median mortality and 95% highest posterior density (HPD) intervals, averaged over sub-line, date, and tube. Background coloured points show observed tube-level mortality percentages; n numbers represent the total number of mosquitoes for that genotype. AA denotes homozygous susceptible (A296/A296), AS heterozygotes (A296/296S), and SS homozygous resistant (296S/296S) mosquitoes. No statistically credible differences between genotypes were detected at this toxic load, as pairwise mortality ORs included 1 (Table 1). The high level of uncertainty (95% HPD intervals) reflects the limited sample size and significant tube-to-tube variability captured by the model.

**Supplementary Figure 4.**
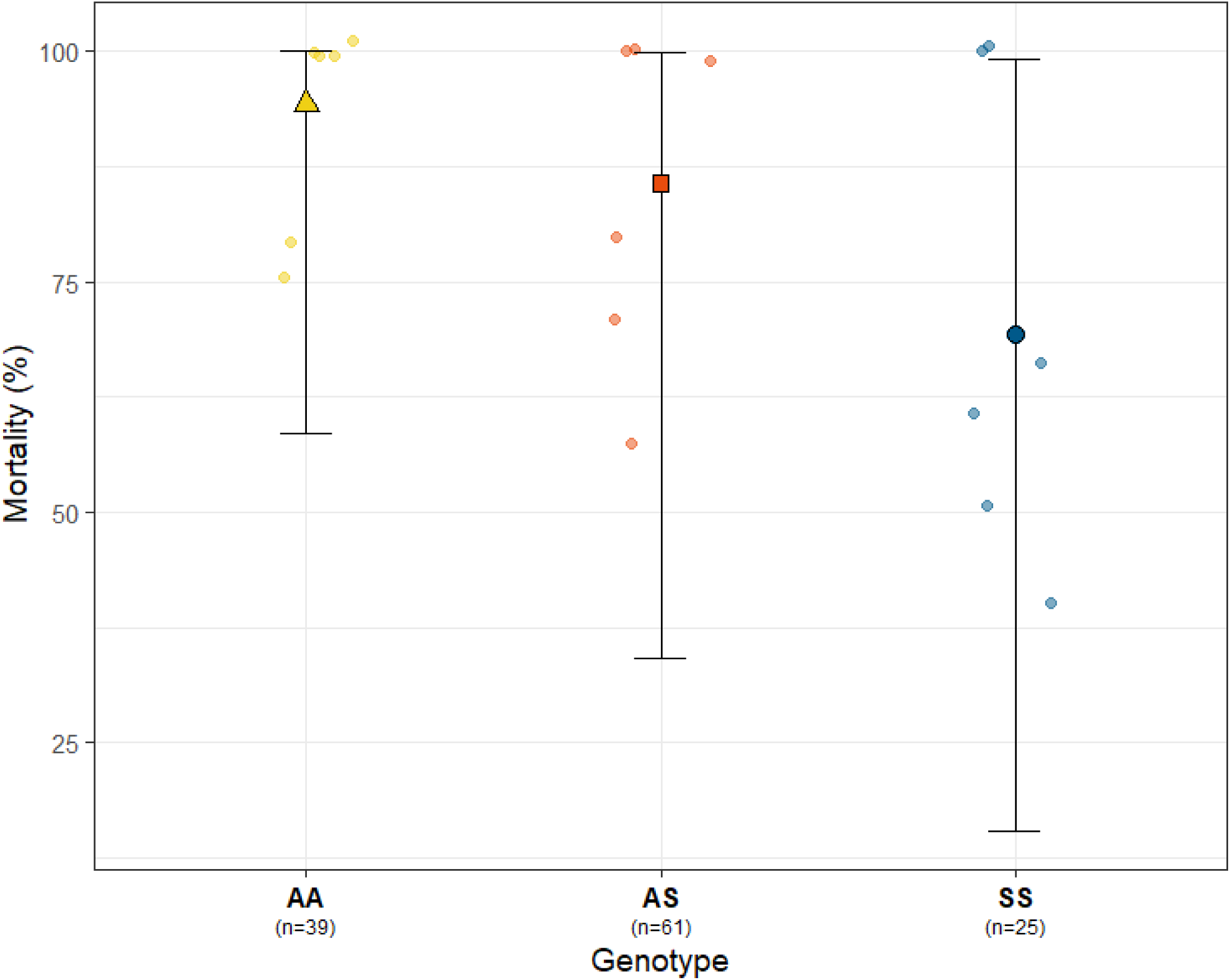
Mortality of adult female *Anopheles funestus* following exposure to 4% DDT for 30 minutes by *Rdl-A296S* genotype. Shapes and error bars indicate model-estimated posterior median mortality and 95% highest posterior density (HPD) intervals, averaged over sub-line, date, and tube. Background coloured points show observed tube-level mortality percentages; n numbers represent the total number of mosquitoes for that genotype. AA denotes homozygous susceptible (A296/A296), AS heterozygotes (A296/296S), and SS homozygous resistant (296S/296S) mosquitoes. No statistically credible differences between genotypes were detected at this toxic load, as pairwise mortality ORs included 1 (Table 1). The high level of uncertainty (95% HPD intervals) reflects the limited sample size and significant tube-to-tube variability captured by the model.

**Supplementary Table 1.** Pooled summary of two independent microinjection sets. Edited events were recovered from one transient female (F4) and from four wild-type females crossed *en masse* to transient males (M6, M12, M25, M33).

| *Embryos Aligned* | *Surviving Larvae* | *G0 Larvae with Transient RFP Expression* | *G0 Transient Individuals Surviving to Adulthood* |  | *Crossing Scheme* | *Produced Larval Offspring* | *Founder Events* |
|---|---|---|---|---|---|---|---|
| 920 | 227 | 100 | **Male** | 25 | X 75 WT-F | 18 | Min. 1<br>Max. 4 |
|  |  |  | **Female** | 27 | X 81 WT-M | 7 | 1 |

**Supplementary Table 2.** Raw mortality data for all conditions tested.

| *Insecticide* | **Exposure time** | **Sub-line** | ***Total tested*** | ***Status*** | ***No.*** | ***Genotype*** |  |  |
|---|---|---|---|---|---|---|---|---|
|  |  |  |  |  |  | **A/A** | **A/S** | **S/S** |
| ***0.4% Dieldrin*** | ***1 hour*** | ***M25*** | 137 | **Alive** | 66 | 0 | 47 | 19 |
|  |  |  |  | **Dead** | 71 | 40 | 29 | 2 |
|  |  | ***M33*** | 303 | **Alive** | 185 | 0 | 141 | 44 |
|  |  |  |  | **Dead** | 118 | 79 | 34 | 5 |
|  |  | ***Pooled*** | 440 | **Alive** | 251 | 0 | 188 | 63 |
|  |  |  |  | **Dead** | 189 | 119 | 63 | 7 |
| ***4% Dieldrin*** | ***1 hour*** | ***M33*** | 57 | **Alive** | 3 | 0 | 0 | 3 |
|  |  |  |  | **Dead** | 54 | 18 | 27 | 9 |
| ***4% Dieldrin*** | ***30 minutes*** | ***M33*** | 235 | **Alive** | 117 | 0 | 63 | 54 |
|  |  |  |  | **Dead** | 118 | 64 | 52 | 2 |
| ***4% DDT*** | ***1 hour*** | ***M33*** | 27 | **Alive** | 0 | 0 | 0 | 0 |
|  |  |  |  | **Dead** | 27 | 6 | 17 | 4 |
| ***4% DDT*** | ***30 minutes*** | ***M33*** | 125 | **Alive** | 20 | 2 | 10 | 8 |
|  |  |  |  | **Dead** | 105 | 37 | 51 | 17 |
| ***0.75% Permethrin*** | ***1 hour*** | ***M33*** | 72 | **Alive** | 0 | 0 | 0 | 0 |
|  |  |  |  | **Dead** | 72 | 20 | 41 | 11 |
| ***0.75% Permethrin*** | ***30 minutes*** | ***M33*** | 20 | **Alive** | 0 | 0 | 0 | 0 |
|  |  |  |  | **Dead** | 20 | 5 | 13 | 2 |
| ***0.05% Deltamethrin*** | ***1 hour*** | ***M33*** | 49 | **Alive** | 0 | 0 | 0 | 0 |
|  |  |  |  | **Dead** | 49 | 9 | 28 | 12 |
| ***0.05% Deltamethrin*** | ***30 minutes*** | ***M33*** | 21 | **Alive** | 0 | 0 | 0 | 0 |
|  |  |  |  | **Dead** | 21 | 5 | 11 | 5 |

**Supplementary Table 3.** Larval development summary statistics.

| ***Sex*** | ***Genotype*** | ***Number*** | ***Median development time, days*** *[IQR]* |
|---|---|---|---|
| **All** | **All** | 482 | **12** [11, 12] |
| **Males** | **All** | 239 | **11** [11, 12] |
| **Females** | **All** | 243 | **12** [11, 12] |
| **All** | **FANG-S/S** | 136 | **12** [11, 13] |
| **All** | **FANG-A/S** | 241 | **11** [11, 12] |
| **All** | **FANG-A/A** | 105 | **11** [11, 12] |

**Supplementary Table 4.** Cox Proportional Hazards regression analysis of larval development time (time-to-pupation). The homozygous susceptible (FANG-A/A) genotype and the female sex were utilised as reference levels. A Hazard Ratio (HR) < 1 (for genotype) indicates a lower probability of pupating at any given time point compared to the reference level, corresponding to a developmental delay (fitness cost). Conversely, an HR > 1 for sex indicates a higher probability of earlier pupation (in this instance indicative of protandry). Significance cutoff: ns P > 0.1;. P < 0.1; * P < 0.05; ** P < 0.01, *** P < 0.001, **** P < 0.0001.

| *Predictor variable (Level)* | *Reference level* | *Coef.* | *Hazard Ratio [95% CI]* | *z* | *P-value* | *Sig.* |
|---|---|---|---|---|---|---|
| **Sex (Male)** | Sex (Female) | 0.11 | **1.12** [0.93, 1.34] | 1.192 | 0.2331 | ns |
| **Genotype (A/S)** | Genotype (A/A) | -0.03 | **0.97** [0.77, 1.23] | -0.226 | 0.8211 | ns |
| **Genotype (S/S)** | Genotype (A/A) | -0.30 | **0.74** [0.57, 0.95] | -2.317 | 0.0205 | * |

**Supplementary Table 5.**
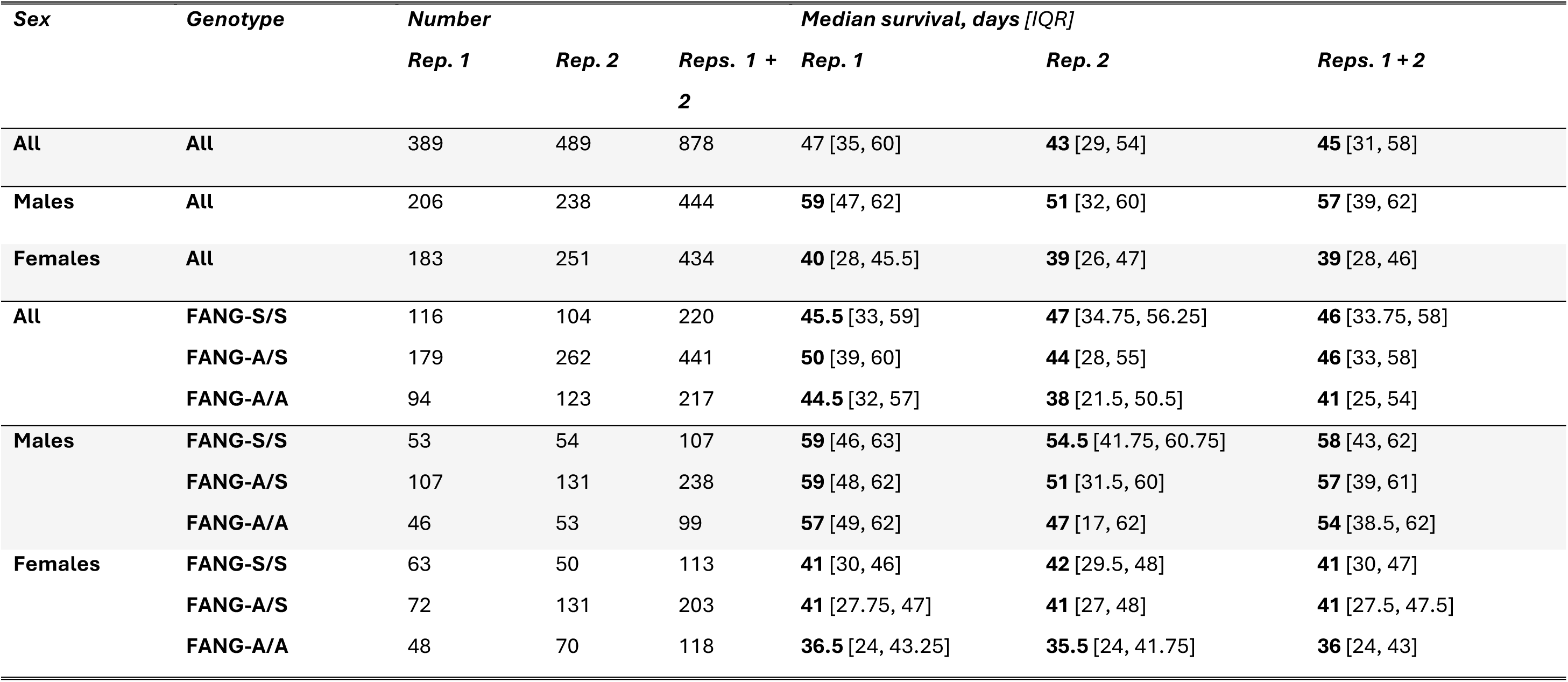
Adult longevity summary statistics.

| ***Sex*** | ***Genotype*** | ***Number*** |  |  | ***Median survival, days*** *[IQR]* |  |  |
|---|---|---|---|---|---|---|---|
|  |  |  |  |  |  |  |  |
|---|---|---|---|---|---|---|---|
| **All** | **All** | 389 | 489 | 878 | 47 [35, 60] | **43** [29, 54] | **45** [31, 58] |
| **Males** | **All** | 206 | 238 | 444 | **59** [47, 62] | **51** [32, 60] | **57** [39, 62] |
| **Females** | **All** | 183 | 251 | 434 | **40** [28, 45.5] | **39** [26, 47] | **39** [28, 46] |
| **All** | **FANG-S/S** | 116 | 104 | 220 | **45.5** [33, 59] | **47** [34.75, 56.25] | **46** [33.75, 58] |
|  | **FANG-A/S** | 179 | 262 | 441 | **50** [39, 60] | **44** [28, 55] | **46** [33, 58] |
|  | **FANG-A/A** | 94 | 123 | 217 | **44.5** [32, 57] | **38** [21.5, 50.5] | **41** [25, 54] |
| **Males** | **FANG-S/S** | 53 | 54 | 107 | **59** [46, 63] | **54.5** [41.75, 60.75] | **58** [43, 62] |
|  | **FANG-A/S** | 107 | 131 | 238 | **59** [48, 62] | **51** [31.5, 60] | **57** [39, 61] |
|  | **FANG-A/A** | 46 | 53 | 99 | **57** [49, 62] | **47** [17, 62] | **54** [38.5, 62] |
| **Females** | **FANG-S/S** | 63 | 50 | 113 | **41** [30, 46] | **42** [29.5, 48] | **41** [30, 47] |
|  | **FANG-A/S** | 72 | 131 | 203 | **41** [27.75, 47] | **41** [27, 48] | **41** [27.5, 47.5] |
|  | **FANG-A/A** | 48 | 70 | 118 | **36.5** [24, 43.25] | **35.5** [24, 41.75] | **36** [24, 43] |

**Supplementary Table 6.** Mixed-effects Cox Proportional Hazards model analysis of adult longevity. The model evaluates the impact of sex and genotype on the mortality risk of *Anopheles funestus* (n = 878) containing the *Rdl*-A296S mutation. Fixed effects include Sex and Genotype, while Replicate was incorporated as a random intercept to account for inter-experimental variation. Reference levels for the analysis are Sex (Female) and Genotype (A/A; susceptible). A Hazard Ratio (HR) < 1 indicates a reduced risk of death (or increased longevity) relative to the reference level, whereas an HR of > 1 indicates an increased risk of death. Schoenfeld residual diagnostics indicated evidence of non-proportional hazards for sex and genotype; hazard ratios should therefore be interpreted as overall summaries of survival differences rather than constant effects over time. Significance cutoff: ns P > 0.1;. P < 0.1; * P < 0.05; ** P < 0.01, *** P < 0.001, **** P < 0.0001.

| *Predictor variable (Level)* | *Reference level* | *Coef.* | *Hazard Ratio [95% CI]* | *z* | *P-value* | *Sig.* |
|---|---|---|---|---|---|---|
| **Sex (Male)** | Sex (Female) | -1.21 | **0.30** [0.26, 0.35] | -15.40 | < 0.0001 | **** |
| **Genotype (A/S)** | Genotype (A/A) | -0.19 | **0.83** [0.70, 0.97] | -2.28 | 0.023 | * |
| **Genotype (S/S)** | Genotype (A/A) | -0.13 | **0.87** [0.72, 1.06] | -1.39 | 0.163 | ns |

**Supplementary Table 7.**
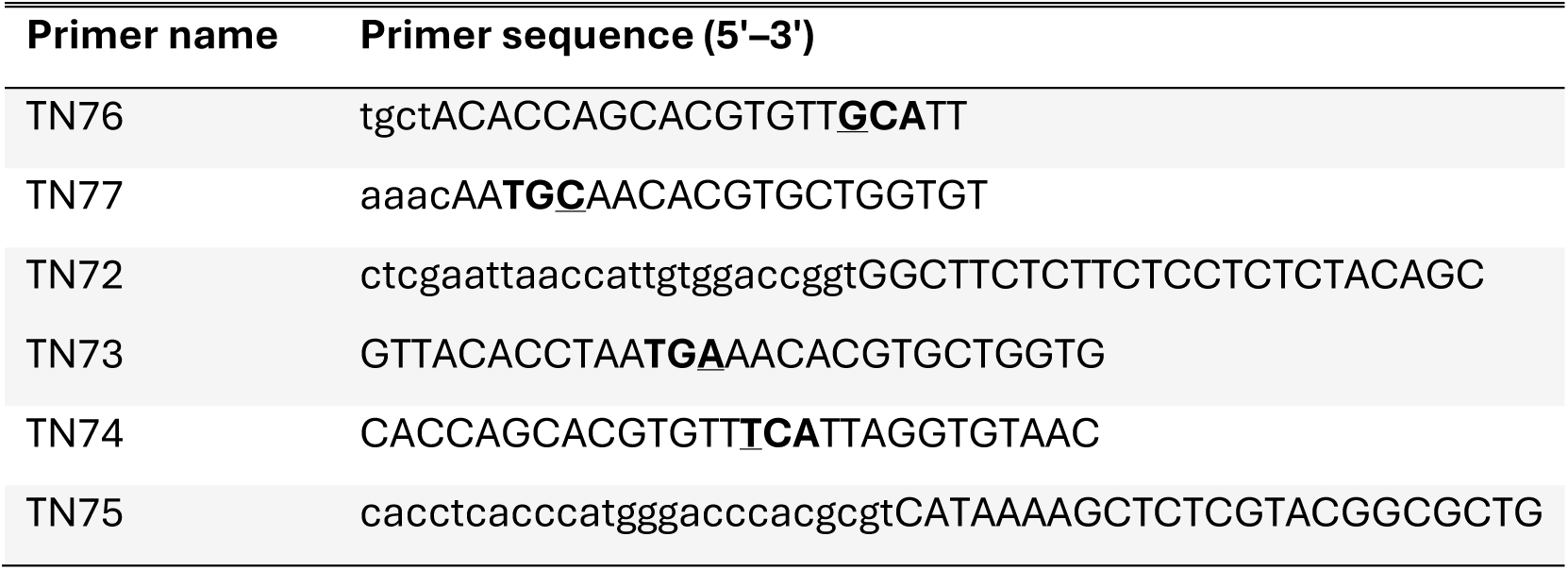
Primers used for cloning the helper plasmid (p68; RdlA296S^Vasa2^) and donor plasmid (p67; RdlA296S-HDR). Bold bases indicate the codon corresponding to alanine in the target sequence incorporated into the helper plasmid (TN76/TN77), or the substituted codon encoding serine in the donor plasmid (TN73/TN74). Underlined bases indicate the nucleotide substitutions introduced to generate the A296S mutation.

**Supplementary Table 8.**
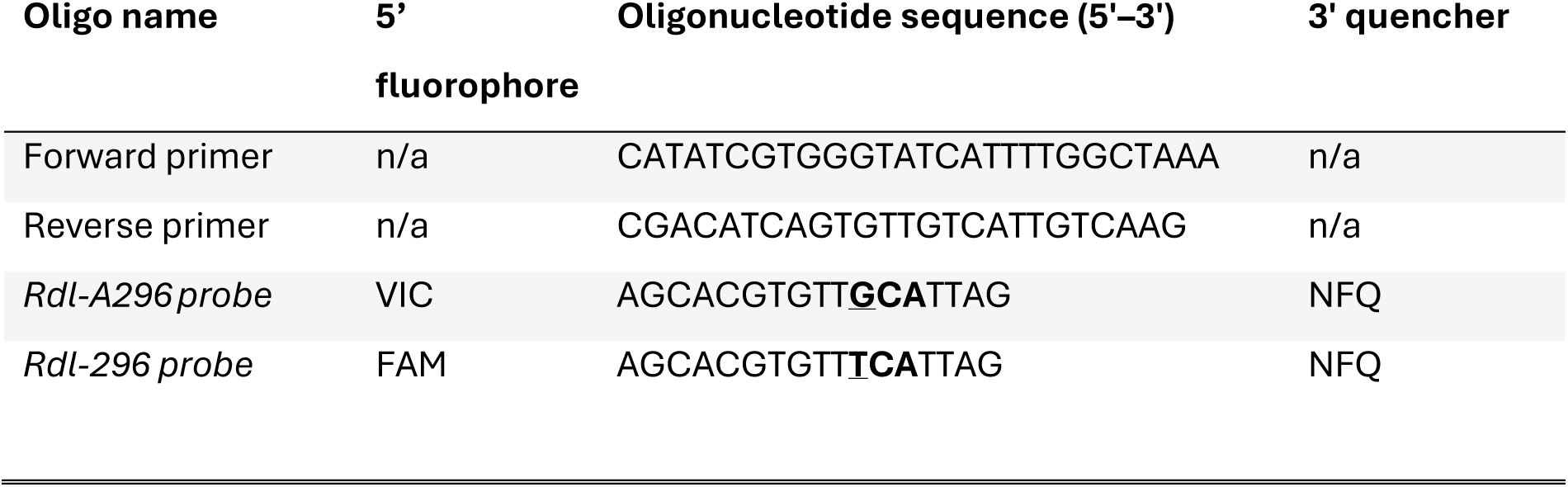
Oligonucleotides used for TaqMan™ genotyping of the Rdl-A296S mutation. The codon corresponding to amino acid position 296 is shown in bold, and the allele-specific nucleotide is underlined.

